# Upper extremity exomuscle for shoulder abduction support

**DOI:** 10.1101/2020.04.25.061846

**Authors:** Cole Simpson, Bryce Huerta, Sean Sketch, Maarten Lansberg, Elliot Hawkes, Allison Okamura

## Abstract

Assistive devices may aid motor recovery after a stroke, but access to such devices is limited. Exosuits can aid human movement and may be more accessible alternatives to current devices, but we know little about how they might assist post-stroke upper extremity movement. Here, we designed an exosuit actuator to support shoulder abduction, which we call an “exomuscle” and is based on a form of growing robot called a pneumatic-reel actuator. We also assembled a ceiling-mounted support for a positive control. We verified that both supports reduce the activity of shoulder abductor muscles and do not impede range of motion in healthy participants (n=4). Then, we measured reachable workspace area in stroke survivors (n=6) with both supports and without support. Our exomuscle increased workspace area in four participants (180±90 cm^2^) while the ceiling support increased workspace area in five participants (792±540 cm^2^). Design decisions that reduce exomuscle complexity, such as leaving the forearm free, likely contribute to performance differences between the two supports. Heterogeneity amongst stroke survivors’ abilities likely contribute to high variability in our results. Though both supports performed similarly for healthy participants, performance differences in stroke survivors highlight the need to validate assistive devices in the target population.

## I. Introduction

Stroke, the death of brain tissue due to interrupted blood supply, is the leading cause of adult disability **Mozaffarian2015** While the typical stroke survivor regains about 70% of lost motor function **prabhakaran2008inter, zarahn2011prediction, winters2015generalizability** stroke remains the most common cause of adult disability in the United States **ovbiagele2011stroke** Due to aging populations and changing lifestyles, the incidence and impact of stroke is expected to increase. Post-stroke permanent disability might be reduced by increasing therapy dosage **mccabe2015comparison, lang2015dose** but rigorously testing large dosages in humans is difficult due to the commitment required of both therapist and patient and the likelihood of interfering with standard care. Assistive devices may help patients to perform more and longer therapies with less therapist involvement.

Assistive devices can be designed to aid patients in a number of ways; one promising approach is to compensate for the effects of gravity. Compensating for gravity has been shown to increase stroke survivors’ reachable workspace **sukal2007shoulder** theoretically increasing the number of tasks that can be performed. The relationship between gravity compensation and increased reachable workspace is attributed to abnormal flexor synergy activation, in which stroke survivors lose the ability to independently control the shoulder abductor and elbow flexor muscles **cheung2012muscle, dewald2001abnormal** Unloading the shoulder abductor muscles by providing gravity compensation reduces activity in the elbow flexor muscles, increasing elbow range of motion **beer2007impact** This gravity compensation strategy has been used in an eight-week therapy in which support was gradually removed, and motor improvements were found **ellis2009progressive, ellis2007act** Because of the simplicity and effectiveness of this approach, gravity compensation or shoulder abduction support is the goal of many assistive devices.

Exoskeletal robots, which use motors and rigid linkages to transmit forces to the user, have been used extensively in research applications **ellis2007act, sukal2007shoulder** However, widespread adoption of rehabilitation exoskeletal robots by practicing healthcare providers has not yet taken place. Some commercially produced passive exoskeletal devices, which use stored potential energy to compensate for gravity, are used in the clinic. Examples of such devices include the Hocoma ArmeoSpring and the SaeboMAS. These commercial devices are rarely used outside the clinic. One key factor limiting adoption may be that existing devices are too expensive and complex to operate. Much cost and complexity stems from safety considerations such as aligning robot and operator joints **schiele2009influence** compensating for the additional mass of the robot, and ensuring stable interactions with users **duchaine2008investigation, kazerooni1995case** These devices are limited by the workspace of the device, which is usually grounded to a table, the floor, or a wheelchair. Wearable devices, grounded to the body instead of the environment, may diversify the contexts in which assistive devices can used.

Soft exosuits can address many challenges associated with traditional exoskeletal robots. While exoskeletal robots use rigid linkages to generate reaction forces, exosuits forgo rigid linkages, relying instead on the user’s skeleton for that purpose **asbeck2014stronger** Doing so removes components and therefore mass from the system, allowing exosuits to be constructed of primarily compliant materials that are lightweight, inexpensive, and require less precise manufacturing. While early exosuit designs focused on assisting with locomotion **asbeck2014stronger, lee2017reducing, awad2017soft** several recent examples actuate the upper extremity **polygerinos2013towards, realmuto2019robotic** Specifically, exosuits have been created to assist with arm movements in stroke survivors **simpson2017exomuscle, oguntosin2015development, natividad2016development, oneill2017soft, lessard2018soft, natividad2018exosleeve, tiseni2019edge** In particular, several of these studies have examined how exosuit use affects muscular activity in healthy users and whether the exosuit limits the user’s range of motion **natividad2016development, simpson2017exomuscle, oneill2017soft, tiseni2019edge** Only one recent study has examined the effects of an upper extremity exosuit on stroke survivors. In that study, exosuit use increased shoulder abduction and flexion range of motion, the degrees of freedom supported by their exosuit, in five stroke survivors **oneill2020inflatable** O’Neill et al. **oneill2020inflatable** do not investigate whether unsupported degrees of freedom, such as the elbow, benefit from exosuit assistance of shoulder abduction support as suggested by previous studies of gravity compensation after stroke. No validations in healthy users were reported in that study.

Here, we design and build a novel inflatable actuator for shoulder abduction support, analogous to gravity compensation (Fig. 1A). We show in healthy participants that the device does not impede range of motion and supports the anterior and medial components of the deltoid muscles in abducting the shoulder, previously shown to be related to post-stroke reachable workspace. We then test in stroke survivors whether exomuscle assistance increases reachable workspace area in the transverse plane (orthogonal to exomuscle support), and whether a ten-minute assisted therapy session changes reachable workspace area. We perform the same set of evaluations using a ceiling-mounted arm support for a positive control. We show an increase in reachable workspace area in the transverse plane in 4 out of 6 stroke survivors. Exomuscle support correlated with increased elbow range of motion, which is not supported by our exomuscle. Although the ceiling-mounted support and exomuscle produced the same results in healthy participants, we show that they produce very different outcomes in stroke survivors, highlighting the need to validate assistive devices in the target population.

**Fig. 1.**
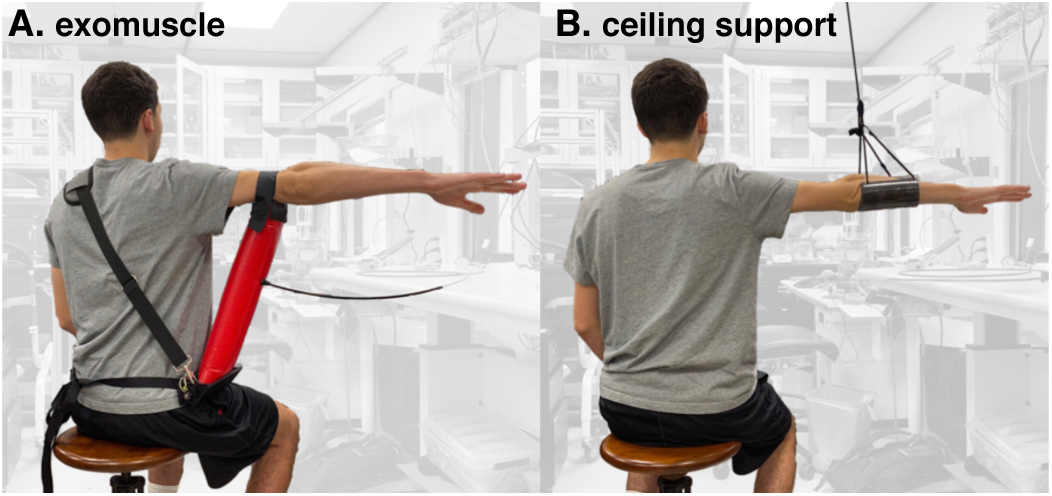
User with (A) an exomuscle and (B) a ceiling support. **A**. The exomuscle is the experimental device in this study. Its design is based on a type of growing robot **hawkes2017soft** called a pneumatic-reel actuator **hammond2017pneumatic B**. The ceiling support is used as a positive control in this study.

## II. Shoulder abduction support device design

Complexity, weight, cost, and safety considerations all limit access to robot-aided therapies. Our goal is therefore to create a simple, lightweight, inexpensive, wearable device capable of aiding in stroke rehabilitation by supporting shoulder abduction. We also constructed a ceiling-mounted support for use as a positive control. These two devices are shown in Fig. 1.

### A. Exomuscle design – experimental device

Exosuits are frequently made using multiple actuators that apply compressive loads on the user’s joints **wehner2013lightweight, mao2011cable, brackbill2009dynamics** However, in early pilot testing, we determined that the forces required to fully support the weight of the arm without standoffs resulted in uncomfortably high compressive loads on the shoulder. Rather than adding standoffs that might catch on objects in the environment to reduce these loads, we designed our device to push the arm from beneath rather than pull from above. Additionally, because our device comprises a single actuator rather than a suit of actuators, we refer to it as an “exomuscle” rather than an “exosuit.”

In Simpson et al. **simpson2017exomuscle** we described a compliant inexpensive wearable exomuscle with a single controlled degree of freedom that assists shoulder abduction in healthy users without restricting motion in other joints. That device controls shoulder abduction angle by modulating the air pressure in a bladder sewn with a hinge-like seam to a chest harness and resting under the arm. That device is effective at lifting the arm and supporting movement while pressurized, but can buckle at small shoulder abduction angles/low air pressures. Here we describe a new actuator that uncouples air pressure and shoulder abduction angle.

Our updated design, based on the pneumatic reel actuators developed by Hammond et al. **hammond2017pneumatic** consists of a plastic bladder reinforced by a fabric tube sewn out of thermoplastic polyurethane (TPU)-coated polyester cloth. The fabric reinforced tube is then wrapped around an axle mounted to the base assembly shown in Fig. 2. In this study, we use a locking disk and pin to fix the shoulder abduction angle so that the elbow rests at shoulder height. For active length control, the lock disk can be replaced with a transmission to a motor as described in Hammond et al. **hammond2017pneumatic** The base assembly is held in place on the wearer’s side by a waist strap and a contralateral shoulder strap, which supports the vertical forces required to support the arm. Our prototype base is made out of polyoxymethylene plastic. The supported arm is held in place on top of the inflated bladder by a hook-and-loop fastener strap. A flow chart of the construction process is shown in Fig. 2.

**Fig. 2.**
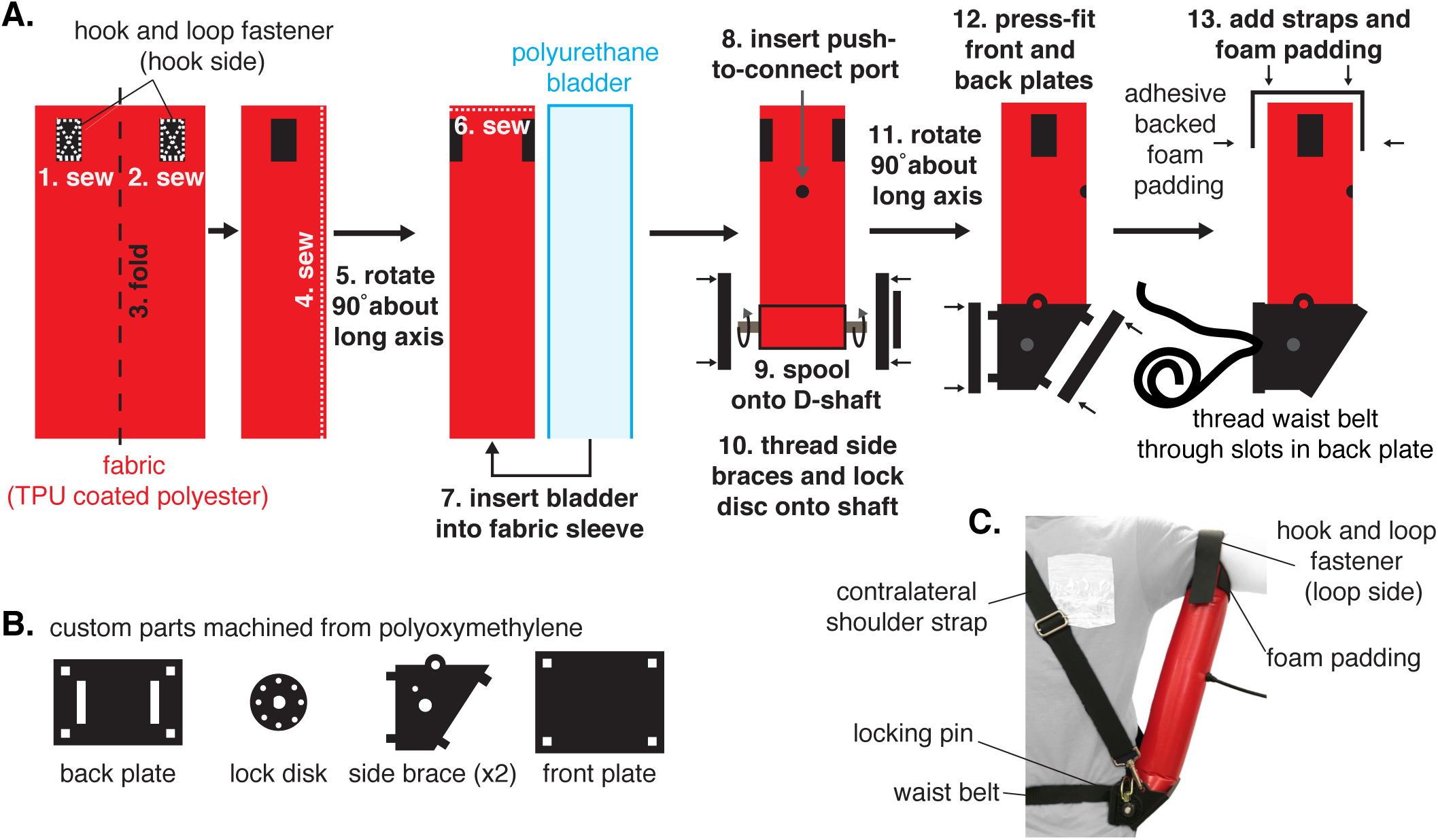
Exomuscle design. **A**. The pneumatic reel exomuscle consists of two main parts, a long bladder and a base that contains the reel. The bladder is constructed of a single piece of TPU coated polyester with hook and loop fasteners sewn near the tip to hold the strap that will later hold the arm in place. The fabric is then sewn into a tube and then twisted one quarter turn around the long axis such that the hook and loop patches are on the sides. The end of the tube is then sewn shut and a polyurethane bladder inserted. A push-to-connect port is then inserted through the first layer of fabric and one side of the bladder to allow for inflation. **B**. The base station is constructed out of four custom parts machined from polyoxymethylene, a back plate with slots for a waist belt, two side braces that include holes for a shoulder strap to attach, a front plate, and a disk brake. The bladder is spooled onto a D-shaft, and the side braces and disk brake are inserted onto the ends of the D-shaft. Shaft collars can be added for extra stability. The back and front plates are then press fit onto the notches on the side braces. Then foam padding and straps can be added as desired. **C**. When completed, the exomuscle base rests against the wearer’s side, supported by the contralateral shoulder strap and waist belt, and underneath the upper arm, supported by another strap.

When completed, this exomuscle has convenient properties for interfacing with human operators. The height of the support can be adjusted without changing the air pressure by spooling and unspooling the bladder on the axle. The arm rests on the inflated tube, which is compliant enough at operational air pressures of 25-50 kPa to provide a comfortable support. Additionally, extra material at the top and bottom of the inflated tube where it transitions from a rectangular cross section (at the ends where the tube is rolled about the axle or sewn with a straight seam) to a circular cross section creates pivot points that act as ball joints. These natural pivots combine with soft tissue compliance from the wearer resulting in a flexible support. Because the exosuit should not interfere with other joints, we attach the exomuscle under the upper arm to avoid crossing the elbow joint, which would require additional compliance about the long axis of the bladder to preserve full elbow range of motion.

### B. Ceiling support – positive control

To examine the effectiveness of the wearable exomuscle design, we need a good-as-possible support to serve as a positive control for comparison. Previous studies examining the effects of gravity compensation in stroke use robotic exoskeletons **ellis2009progressive** or low-friction air sleds to support the arm **beer2007impact** In both cases, the workspace of the exoskeleton or table supporting the air sled requires that the workspace task be broken into segments with the robot or table repositioned in the middle to complete the workspace measurement. These devices also add inertia to the arm, a potential confound.

Instead, we constructed a low-inertia, large workspace sling support **stienen2009freebal, essers2013inverse** We suspended an arm rest from the ceiling using 165 cm of high-strength nylon rope. The arm rest was placed under the forearm as close as possible to the center of mass of the arm for each participant. We positioned each participant such that their elbow was directly beneath the attachment to the ceiling.

## III. Device effects in unimpaired population

Before testing in a vulnerable population such as stroke survivors, we wanted to ensure that the exomuscle produces the intended effects and test for confounding effects in unimpaired participants. To that end, we recruited four participants with no indicated musculoskeletal deficits or injuries (3 males, 1 female; age: 26±4 yrs; height: 180±6 cm; mass: 72±10 kg; all right-hand dominant). Participants gave their informed consent in accordance with the policies of the Stanford Internal Review Board. We tested if the exomuscle was able to offload the effort of the shoulder abductor muscles most commonly affiliated with abnormal flexor synergy activation. We also conducted a test to examine whether the exomuscle limits the range of motion of healthy users.

### A. Deltoid muscular effort

To determine if our exomuscle assists the user’s muscles with shoulder abduction, we measured muscle activity in the shoulder muscles related to post-stroke workspace impairments while participants performed isometric and dynamic tasks. We measured muscle activity from the anterior and medial components of the deltoid muscle using a Delsys Bagnoli 2 EMG system (Natick, MA, USA) read with a National Instruments USB-600X DAQ (Austin, TX, USA) at 1000 Hz. In the isometric tasks, participants were asked to hold their right arm at the height of the shoulder (90° shoulder abduction) with elbow fully extended for three seconds in each of three postures: (1) hand directly in front of the shoulder (90° shoulder flexion), (2) hand directly out to the side (0° shoulder flexion), and (3) halfway in between those postures (45° shoulder flexion). In the dynamic tasks, participants were asked to start in an initial posture in which the arm is raised to the height of the shoulder (90° shoulder abduction), the elbow fully flexed, and shoulder rotated such that the hand is directly in front of the shoulder. Participants were then asked to reach to each of the three target postures described in the isometric tasks (shoulder abducted 90°, elbow fully extended, shoulder flexed to 90°, 0°, and 45°) before returning to the initial posture. Each task was demonstrated first by the experimenter, participants were asked to confirm they understood the instructions, and the experimenter monitored performance of each task to ensure the correct postures were attained. Each participant completed both isometric and dynamic tasks in three conditions: (1) unassisted, (2) exomuscle support, and (3) support from the ceiling support described in Section II-B.

Electromyograms (EMG) from each muscle were high-pass filtered with a cutoff frequency of 30Hz, rectified, and then low-pass filtered with a cutfoff frequency of 6Hz to create linear envelopes. All filters were 8th order, zero-phase shift Butterworth filters (butter and filtfilt functions in MATLAB, Natick, MA, USA). We then computed the average muscle activation in each trial and muscle and compiled all average muscle activations resulting from a given support condition (unsupported, exomuscle, or ideal) and task condition (isometric or dynamic) into a set. Because a one-sample Kolmogorov-Smirnov test showed that the data in some sets were not normally distributed, we used the non-parametric Kruskal-Wallis test followed by Tukey’s test to compare muscle activation across support conditions.

Exomuscle use reduced muscle activity by 79.5% (*p* = 1.7 × 10^−9^) and 58.6% (*p* = 9.6× 10^−6^) on average in the isometric and dynamic tasks, respectively (Fig. 3). The ideal support reduced muscle activity by 83.5% (*p* = 6.1 × 10^−11^) and 52.9% (*p* = 4.6 × 10^−7^) on average in the isometric and dynamic tasks, respectively (Fig. 3). The average muscle activity during exomuscle use was not significantly different from that measured while using the ideal support use in either isometric or dynamic tasks with an *α* = 0.05 confidence level. Both assistive devices support shoulder abduction as intended.

**Fig. 3.**
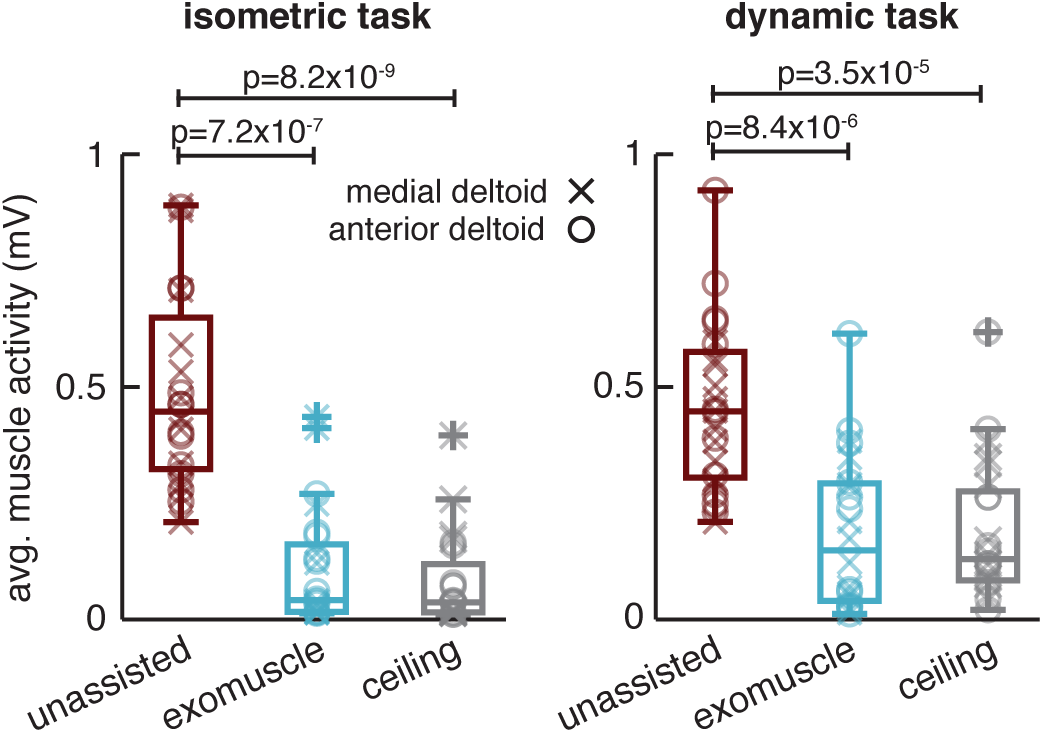
Healthy participant muscular effort in different support conditions (n=4). The average anterior and medial deltoid muscle activations were computed for each trial of the isometric and dynamic tasks. These were combined into sets according to support type: unassisted, exomuscle, or ceiling support. In both isometric and dynamic tasks, muscle activity decreased with both exomuscle and ceiling support use. There was no difference between the muscle activations recorded in the two support conditions at an *α* = 0.05 confidence level in either task.

### B. Range of motion

To be useful in a wide range of tasks, the exomuscle should not impede the motion of the upper extremity while supporting shoulder abduction. To measure upper extremity range of motion, we used motion capture (Impulse X2E, Phasespace, San Leandro, CA, USA) to measure body kinematics at 135 Hz while participants were instructed to move to the extrema of elbow flexion and shoulder flexion with no assistance, with exomuscle support, and with the ceiling support. We also assessed shoulder rotation with no assistance and with exomuscle support. We did not test shoulder rotation with the ceiling support because it attaches at the forearm and thus does not allow shoulder rotation.

Motion capture data was low-pass filtered at 20 Hz using an 8th order, zero-phase shift Butterworth filter (butter and filtfilt functions in MATLAB, Natick, MA, USA). Any gaps in the marker trajectories caused by temporary occlusions of the marker were filled using cubic interpolation (interp1 function in MATLAB). We computed the elbow angle using the angle between the forearm vector (vector connecting the LED markers placed on the lateral humeral epicondyle to the ulnar epicondyle) and upper arm vector (vector from the acromion to the lateral humeral epicondyle). We computed the shoulder flexion angle as the angle between the trunk vector (vector connecting the acromion with the C7 vertebrate) and the upper arm vector. We computed the shoulder internal/external rotation angle as the angle between the forearm vector and the vertical axis of the motion capture coordinate frame. For comparison across conditions, we compute the range of motion as the difference between each extrema. For example, the elbow range of motion is the difference between the peak elbow flexion angle and the peak elbow extension angle. We then grouped the ranges of motion for elbow flexion/extension and shoulder flexion/extension into a single set for each support condition. Because a one-sample Kolmogorov-Smirnov test showed that the data in some of these sets were not normally distributed, we used the non-parametric Kruskal-Wallis test to compare range of motion across support conditions. We used the non-parametric Wilcoxon signed-rank test to compare shoulder internal/external rotation in the unassisted and exomuscle support conditions because the ceiling support does not allow for this motion.

The results of the range of motion analysis are shown in Fig. 4. Neither the exomuscle nor ceiling support changed the range of motion for shoulder flexion/extension or elbow flexion/extension (*p* = 0.97, Kruskal-Wallis). The exomuscle did not reduce the shoulder internal/external rotation range of motion (*p* = 0.89, Wilcoxon signed-rank test). Thus the exomuscle and ceiling support do not impede the motion of the unsupported joints.

**Fig. 4.**
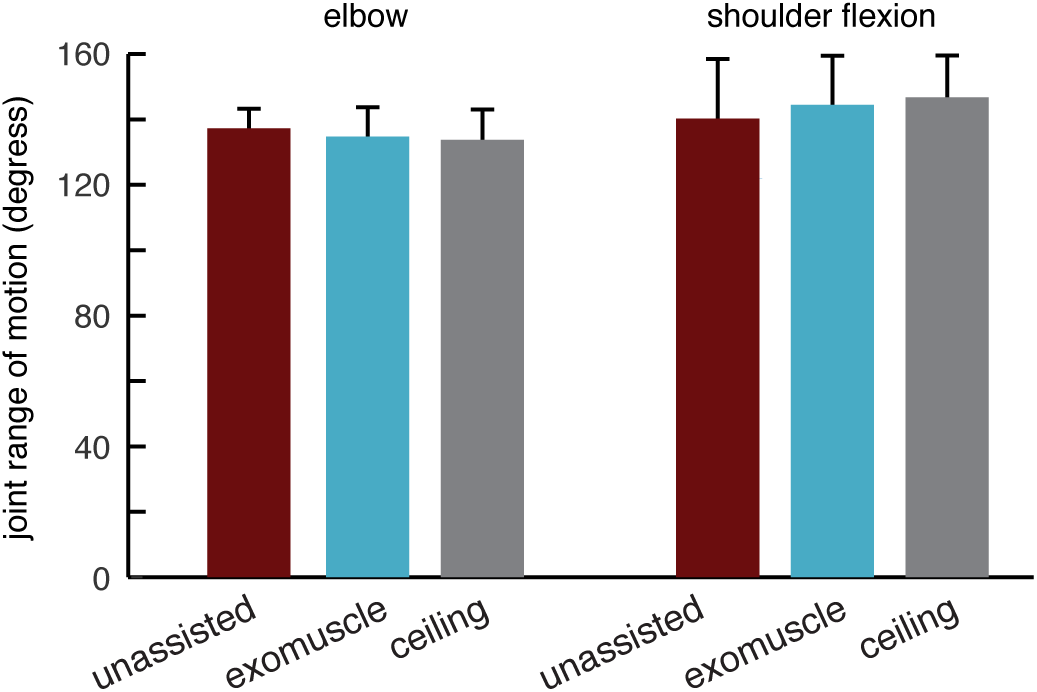
Healthy participant range of motion in different support conditions (n=4). In addition to supporting shoulder abduction, the exomuscle is designed to minimize its impact on other upper extremity degrees of freedom. Here, we measured range of motion in the shoulder and elbow in the unassisted case (red), while using an exomuscle (blue), and with support from the ceiling-mounted device (gray). The range of motion did not change across the conditions (*p* = 0.97, Kruskal-Wallis test).

## IV. Device effects on post-stroke motor performance

Our device was designed to support shoulder abduction, previously shown to increase the reachable workspace area in stroke survivors **beer2007impact, sukal2007shoulder** by offloading abnormal flexor synergy effects **cheung2012muscle, dewald2001abnormal** To determine whether the shoulder abduction assistance from our device increases post-stroke reachable workspace as expected, we recruited 6 stroke survivors with upper extremity motor impairments. We screened participants during the recruitment process to include those with arm weakness whose stroke occured more than 6 months before the study date and exclude those with conflating health problems, sensory deficits, or that experience pain during passive movements of the arm. All participants gave their informed consent in accordance with the policies of the Stanford Internal Review Board. We administered the upper extremity portion of the Fugl-Meyer exam and tested for visual and tactile neglect in all participants. We also administered the manual muscle test (MMT) and modified Ashworth Scale (MAS) on the biceps, triceps, and deltoids of each participant. Because of safety concerns in lifting the arm, we also checked all participants for scapulo-humeral rhythm. The results of these assessments as well as demographic information are shown in Table I. We measured the reachable workspace of all participants in 3 support conditions: unassisted, exomuscle support, and ceiling support. We also tested an indicator of flexor synergy activation in each condition and tested for session effects.

### A. Methods

We designed our exomuscle to aid shoulder abduction and thus increase the reachable workspace of post-stroke users while wearing the device. In pilot studies, however, one participant’s unassisted reachable workspace appeared to increase immediately after a short period of exercise with exomuscle support when compared to unassisted reachable workspace before the exercise. Such session effects are rarely reported **mignardot2017multidirectional, huang2012augmented** and the neurophysiological mechanisms for such spontaneous recovery, if they exist, could be useful to investigate. We thus designed a protocol to not only assess (1) reachable workspace and (2) flexor synergy activation in each support condition, but also (3) whether a short period of assisted exercise could produce short-term motor improvements.

Our protocol consists of a set of assessments performed with each of the 3 support conditions. In each assessment, the reachable workspace was measured using the protocol described below in Section IV-A1 and a proxy for flexor synergy activation was measured using the protocol described below in Section IV-A2. Each participant completed the first set of assessments unassisted to get baseline measurements. Participants were then randomly assigned a support condition (exomuscle or ceiling support) and completed a trial with that support condition followed by a trial with the other support condition as shown in Fig. 5. Each trial consists of a workspace and flexor synergy evaluation with the assigned support followed by a ten-minute intervention trial designed to replicate the exercise that appeared to produce a session effect in a pilot study. In this intervention trial, participants performed the same clockwise and counterclockwise circular movements with the hand used to measure the reachable workspace (see Section IV-A1). The direction of the circular movements was randomly determined and repeated for 1 minute. Up to three 1-minute rest periods were allowed as needed. The support was then removed and participants performed another workspace and flexor synergy assessment with no support to test for any effects of the exercise. Participants then rested for ten-minutes before performing another workspace and synergy assessment to determine whether any exercise effects had worn off. Participants were given one more ten-minute rest before a final assessment. Participants then completed the same set of tests with the other support condition, exomuscle or ceiling support.

**TABLE I.** Stroke survivor demographics and clinical assessment scores.

| ID | Age | Sex | Dominant arm | Affected arm | Years since stroke | FMA UE | MAS | MMT | Neglect | Rhythm |
| --- | --- | --- | --- | --- | --- | --- | --- | --- | --- | --- |
| 1 | 48 | M | R | R | 3.5 | 36 | [1+, 1+, 2] | [2, 4, 4] | N/N | Y |
| 2 | 53 | M | R | L | 3 | 25 | [0, 0, 0] | [5, 4, 4] | N/N | Y |
| 3 | 70 | M | R | L | 11 | 14 | [0, 1, 0] | [2-, 2-, 2+] | N/N | Y |
| 4 | 44 | F | R | L | 2 | 21 | [0, 0, 2] | [2-, 4, 2] | N/N | Y |
| 5 | 50 | F | R | L | 1 | 17 | [4, 1+, 2] | [4, 2+, 3+] | N/N | Y |
| 6 | 41 | M | R | L | 20 | 47 | [0, 2, 0] | [4, 4, 5] | N/N | Y |

Abbreviations: M = male, F = female, R = right, L = left, N = no, Y = yes
FMA UE = upper extremity portion of the Fugl-Meyer assessment, MAS = modified Ashworth scale, MMT = manual muscle test
Neglect = visual and tactile neglect, Rhythm = scapulohumeral rhythm.
Scores on the MAS and MMT are presented in the order: [deltoid, biceps, triceps].
Visual and tactile neglect results are presented in the order: visual/tactile.

**Fig. 5.**
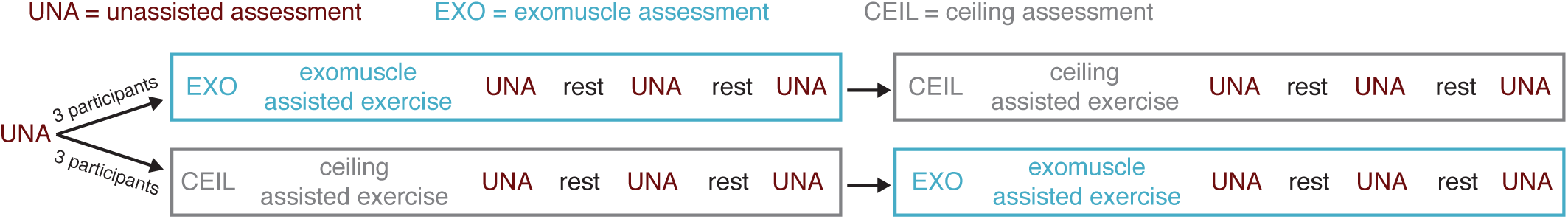
Experimental protocol followed by stroke survivors. Participants performed a baseline assessment without support from either test device. Participants were then randomly assigned to use the exomuscle or ceiling support first. They performed an assessment with that support followed by a 10-minute supported exercise. Follow-up assessments punctuated by 10-minute rests were performed after each exercise to gauge residual effects of device use. Participants then repeated this set of tests with the other support. Each assessment consists of a workspace measurement and the recording of a proxy for flexor synergy activation.

#### 1) Workspace assessment

We followed the protocol used to measure the reachable workspace in stroke survivors from Sukal et al. **sukal2007shoulder** with small modifications to accommodate our devices and equipment. In every case, participants were instructed to individually flex and extend the elbow and shoulder to trace the largest possible circle that they could reach while keeping the hand and elbow raised to the height of the shoulder. In order to record the largest workspace, participants performed 6 circles per measurement, 3 in each direction with the order randomly determined, and were coached by the experimenter who demonstrated the ideal form throughout. In each trial, arm kinematics were recorded using motion capture (Impulse X2E, Phasespace, San Leandro, CA, USA) at 135 Hz with LED markers placed on the C7 vertebrate, acromion, clavicle, olecranon, lateral humeral epicondyle, radial styloid, ulnar styloid, and first knuckle.

Motion capture data was low-pass filtered at 20 Hz using an 8th order, zero-phase shift Butterworth filter (butter.m and filtfilt.m in MATLAB, Natick, MA, USA). Any gaps in the marker trajectories caused by temporary occlusions (less than 0.2 seconds) of the marker were filled using cubic interpolation (interp1 function in MATLAB). Any occlusions lasting longer than 0.2 seconds were ignored in our analyses rather than interpolated to avoid generating incorrect data. We computed the reachable workspace of the hand using the boundary function in Matlab, which computes a concave boundary around the input data and computes the area of that boundary. However, because the original task described by Sukal et al. **sukal2007shoulder** required that the arm remain at the height of the shoulder throughout the experiment, we disregarded datapoints in which the hand (defined as the marker on the ulnar epicondyle) or elbow (defined as the marker on the lateral humeral epicondyle) fell below 20 cm below the height of the shoulder (defined as the marker on the acromion). Due to occlusions in some participants, we substituted the acromion for the C7 marker in all trials. Because the 20 cm threshold selected has the potential to impact the results, we performed a sensitivity analysis of this value presented in the Appendix. We also computed the range of elbow and shoulder flexion/extension achieved during the workspace recordings using the same method described in Section III-B.

#### 2) Flexor synergy assessment

The increase in workspace that normally occurs with gravity compensation is thought to be due to unloading of the flexor synergy. To estimate the relative magnitude of the flexor synergy effect, Ellis et al. **ellis2017flexion** developed an experiment to approximate flexor synergy activation in which the affected arm is placed in a posture that activates the deltoid muscles – and thus the flexor synergy in those that express abnormal flexor synergy – while measuring the activity of the biceps muscles. We copied this experiment with each of our support conditions by asking each participant to hold a posture for three seconds in which the hand and elbow were raised to shoulder height with the hand resting in front of the shoulder and the elbow flexed approximately 70° while the activity of the biceps was measured using a Delsys Bagnoli 2 system (Natick, MA, USA) read with a National Instruments USB-600X DAQ (Austin, TX, USA) at 1000 Hz.

EMG from each participant were high-pass filtered with a cutoff frequency of 30Hz, rectified, and then low-pass filtered with a cutfoff frequency of 6Hz to create linear envelopes. All filters were 8th order, zero-phase shift Butterworth filters (butter and filtfilt functions in MATLAB). We then computed the average muscle activation in each trial.

### B. Results

#### 1) Workspace

Workspace area increased with exomuscle support in 4 out of 6 participants compared with 5 out of 6 with ceiling support (Fig. 6). The two participants who did not increase in workspace with exomuscle support were the two participants with the largest baseline workspace area (S2 and S6, Fig. 6). Only participant 4 did not increase in workspace area with ceiling support. This participant also exhibited the most fatigue and spasticity effects. Note that we present the workspace area change relative both to the baseline measurement and relative to the previous unassisted trial in Fig. 6B. We discuss the comparison against the previous unassisted trial here because it is likely the more accurate measurement due to the effects of fatigue and spasticity discussed in Section IV-B3.

**Fig. 6.**
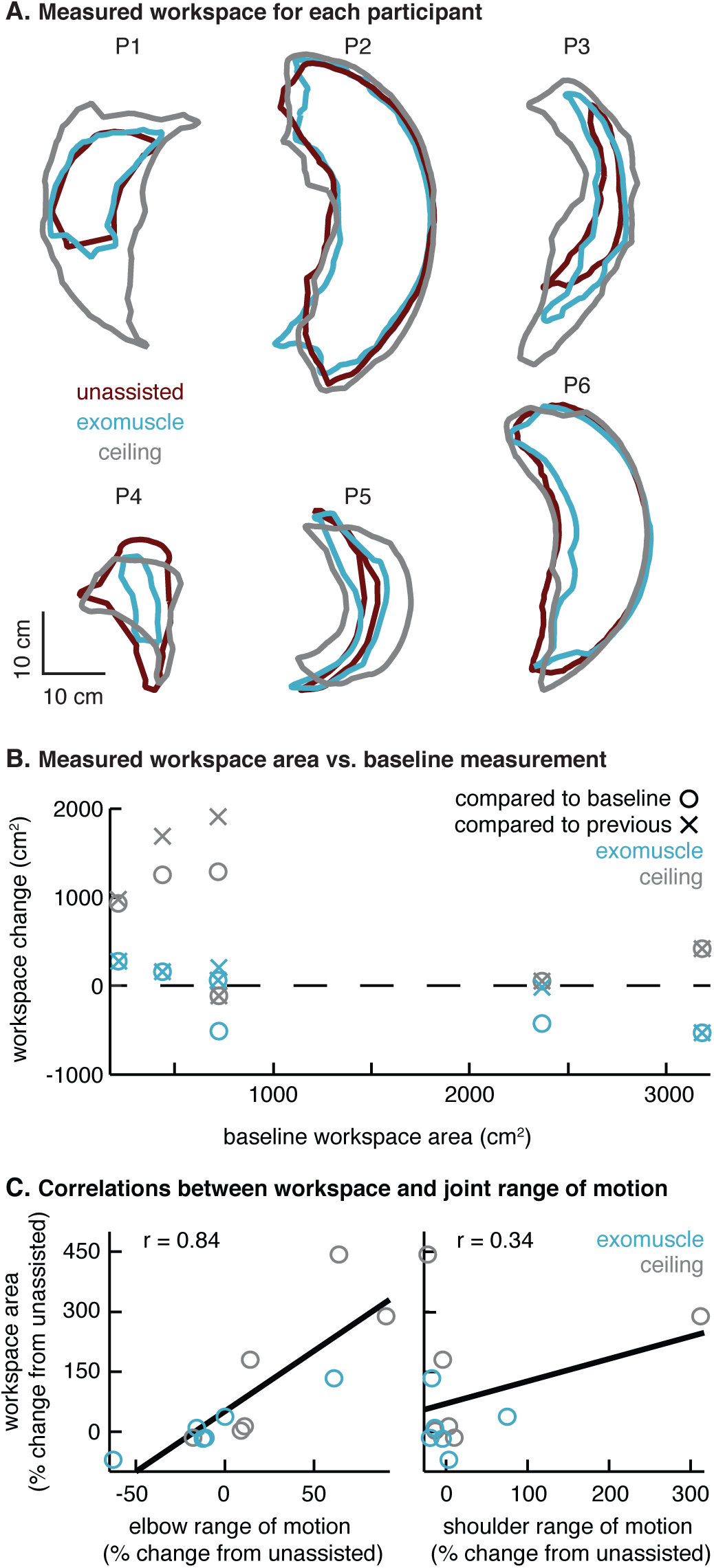
Stroke survivor workspace measurements with different support conditions (n=6). **A**. Workspace area measured in each participant in three support conditions: unassisted, exomuscle, and ceiling support. **B**. Workspace area increased for only three participants using the exomuscle (blue hollow circles) while the ceiling support increased workspace in five participants (gray hollow circles). Pariticpants with small unassisted workspaces received more benefit from both devices. Four participants increased in workspace when compensating for fatigue by comparing against the previous unassisted trial, rather than the first. **C**. Elbow range of motion was strongly correlated with workspace area (r=0.84). This correlation supports the argument that the support devices offload the flexor synergy, which constrains elbow range of motion. Shoulder range of motion was weakly correlated with workspace area (r=0.34).

Changes in workspace area were strongly correlated with changes in elbow range-of-motion (r=0.84), while shoulder range-of-motion was weakly correlated with workspace area (r=0.34, Fig. 6C.), consistent with Sukal et al.’s findings **sukal2007shoulder**

#### 2) Flexor synergy

Biceps activation during a shoulder abduction task, a proxy for flexor synergy activation, decreased in 4/6 and 5/6 participants using the exomuscle and the ceiling support, respectively. Biceps activations were weakly correlated with elbow range of motion (r=0.31) and workspace area (r=0.41) (Fig. 7). Though the correlations shown in Fig. 7 are weak, they show the expected trend: that reductions in flexor synergy activation will lead to increased elbow range of motion and increased reachable workspace. This correlation might be weak due to our limited ability to screen for participants with abnormal flexor synergy activation, adding noise to this measurement.

**Fig. 7.**
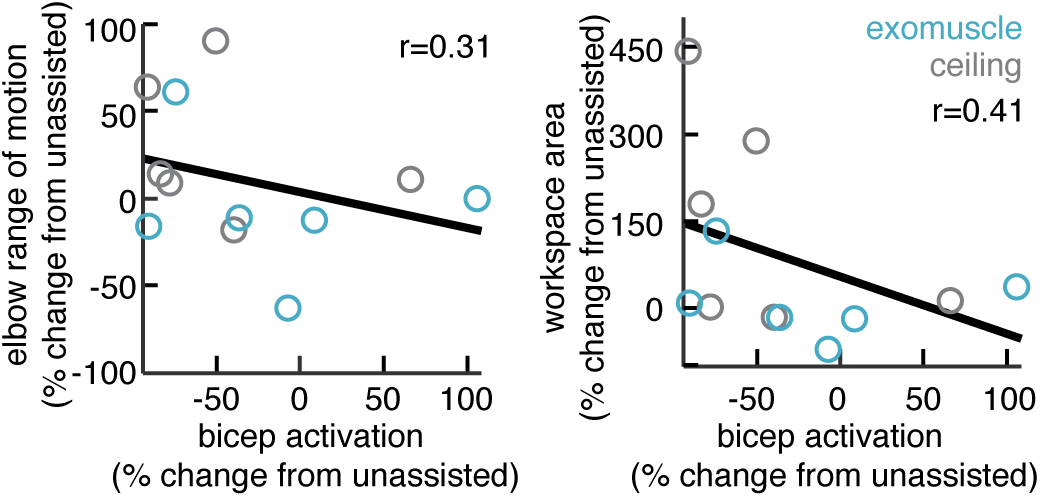
Correlations between biceps activation and elbow range of motion and workspace area. We measured biceps muscle activation as a proxy for flexor synergy activation **ellis2017flexion** Changes in bicep activation were weakly negatively correlated with both elbow range of motion (left) and workspace area (right).

#### 3) Session effects

We did not see single-session increases in workspace area after our intervention. Rather, we saw short-term decreases in workspace related to use of the arm, likely due either to fatigue or use-dependent effects of spasticity (Fig. 8). Workspace area decreased more after exomuscle exercise than after exercise with the ceiling support (p= 0.03, Wilcoxon rank sum), suggesting exomuscle use was more fatiguing than use of the ceiling support. Because participants did not recover these losses in workspace area between the first and third unassisted assessments (p=0.79, Wilcoxon sum rnk), this time-dependent effect is likely to have impacted our results. If we compare workspace area while using the exomuscle to the previous trial in which assistance was not used rather than the baseline measurement, 4/6 participants increase in workspace by an average 180 ±90 cm^2^.

**Fig. 8.**
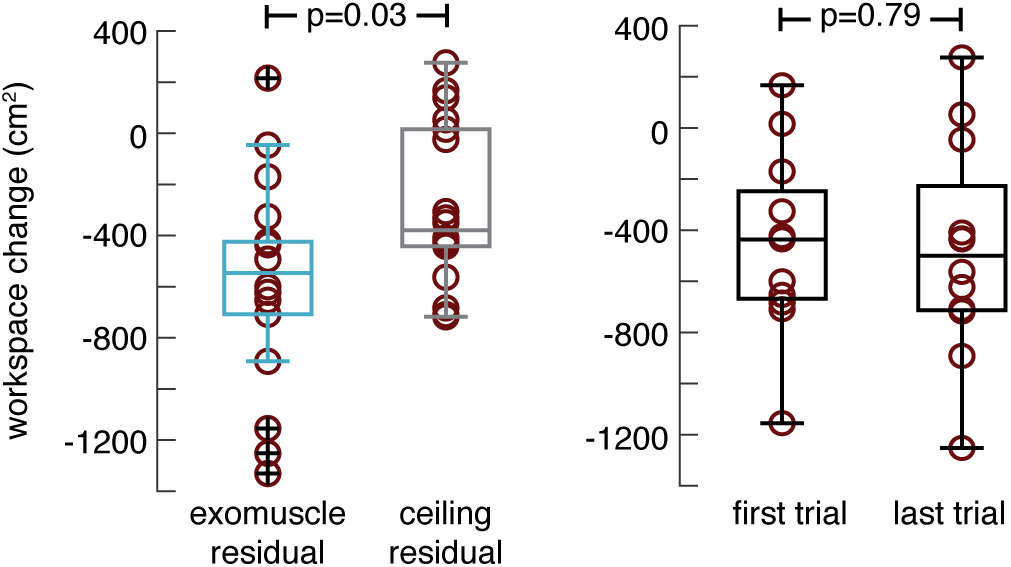
Session effect on unassisted workspace area. Workspace area decreased more after exercise with the exomuscle than with the ceiling support (left, p=0.03, Wilcoxon rank sum test), suggesting that exomuscle use was more fatiguing than the ceiling support. Workspace area did not return to baseline after the two ten-minute rest periods (right, p=0.79, Wilcoxon rank sum test), suggesting that fatigue or spasticity may have played a role in decreasing workspace measurements in the second half of the experiment. To control for this, we compared workspace measurements to the most recent unassisted trial in addition to the baseline trial in Fig. 6.B.

## V. Discussion

We designed a single degree-of-freedom exosuit actuator that we call an exomuscle to support shoulder abduction, verified that it performs as intended in healthy participants, and tested its efficacy in increasing a motor performance metric in stroke survivors. Several recent devices incorporate multiple controlled degrees of freedom to command a larger range of actions in the wearer. We take the opposite approach in attempting to assist only a single degree of freedom shown in other works to be effective in improving a measure of post-stroke motor capability, reachable workspace area. We verified in healthy subjects that our exomuscle reduces the activity of shoulder abductor muscles required for support against gravity and implicated in reduced workspace area. We also verified that the exomuscle does not impede joint range of motion of the shoulder or elbow. We then measured workspace area in six stroke survivors, finding an increase in workspace area in four with exomuscle support. We compared the exomuscle with a ceiling support serving as a positive control, which increased workspace area in five participants. Though the two performed similarly in healthy participants, the ceiling support outperformed the exomuscle in every measure in stroke survivors. This result highlights the need to study device effects in the target population.

Several design factors may contribute to the difference in performance between the two devices in stroke survivors. Certain design concessions that were made to make the exomuscle wearable are not necessary for the ceiling support. The most noticeable of these is that the ceiling support lifts the user’s forearm while the exomuscle supports the wearer’s upper arm, leaving forearm support to the user. This concession enables exomuscle users to internally rotate the shoulder and allows for a simpler design than would be required to support the forearm throughout the full range of shoulder-plus-elbow motion. This concession made little difference to healthy users, who were able to support their own forearms while exploiting the exomuscle for support. Stroke survivors, on the other hand, frequently needed the additional forearm support provided by the ceiling support to maintain the experiment posture.

Stroke survivors have different levels of motor ability depending on the size and location of the lesion and on treatment. Stroke survivors frequently experience a combination of sensorimotor deficits including abnormally coupled joints **Dewald1995, beer2007impact, dewald2001abnormal** increased noise in sensing and estimation **Cusmano2014** muscular weakness **Bourbonnais1989** and spasticity **Watkins2002** Our exomuscle was designed to assist individuals with abnormally coupled joints in the form of the flexor synergy **duysens2013flexion** Our screening procedure, however, likely failed to distinguish between individuals with an abnormal flexor synergy and those with muscular weakness for example. These sensorimotor deficits can have very different effects on motor performance **sketch2017simulating** which might explain in part why some of our participants increased in reachable workspace by 130% while others decreased by up to 70%.

Exosuits, which have no rigid frame, require that the user’s musculoskeletal system must support the additional loads generated by the device. In many cases, these loads are compressive **wehner2013lightweight, mao2011cable, brackbill2009dynamics, lessard2018soft, mao2011cable** However, our device can produce tensile loads on the shoulder complex, which must be supported by soft tissue. Some stroke survivors have difficulty supporting tensile loads in the shoulder, even in unloaded scenarios, resulting in subluxation. Though commercial braces exist to support against subluxation in stroke survivors that could be used in parallel with our device, a better approach might be to design assistive devices that either generate compressive loads on the shoulder in a comfortable range or those that add no net resultant force to the shoulder as in traditional exoskeletons whose frames support their own reaction forces.

Open challenges for this device include active control necessitating transparent intent recognition and onboarding of all actuators and power sources. Simple, inflatable devices such as ours still require an actuator, compressor, power supply, controller, and sensing for transparent untethered operation. Given that our device is meant to compensate for gravity, a predictable force whose effects vary only with body state, purely mechanical devices can be designed to accomplish the same thing with 0 controlled DOFs **herder2005development, cardoso2002conceptual, van2019design, babik2019play** Such mechanisms, can provide shoulder abduction support that varies with posture to negate the effects of gravity on a wearer’s arm without actuators, compressors, power supplies, controllers, or sensing. These mechanisms’ rigid frames react applied loads, meaning that no net force is applied to the shoulder complex that may cause discomfort in stroke survivors.

While our exomuscle is designed for the needs of stroke survivors, other populations might also benefit from similar devices. Gravity supported therapy has shown to improve clinical measures of motor ability in participants with cerebral palsy **elshamy2018efficacy** Gravity-compensating devices can also provide movement benefits to individuals with muscular dystrophy **kooren2015design** though most extant examples are mounted to wheelchairs and are not wearable.

## Appendix A. Sensitivity to selected threshold value

To estimate the workspace area at the height of the shoulder in our experiment, we disregarded datapoints in which the hand or elbow markers dropped below the shoulder marker by threshold of 20 cm (Fig. 6). However, because this threshold has the potential to influence our results, we perform a parameter sweep on the value of the threshold. Our parameter sweep shows that workspace measurements using the ceiling support are insensitive to the selected margin, likely due to the forearm support provided.

## Acknowledgments

We thank Mark Cutkosky for the use of the EMG system and Cara Welker, Cara Nunez, Michael Raitor, Zonghe Chua, and the rest of the Stanford CHARM lab for their constructive comments and prototype design testing. We thank Marion Buckwalter, Julie Muccini, Kara Flavin, Elizabeth Osborn, Esther Rah, Emily Huang, Michael Sharp, and the rest of the Stroke Collaborative Action Network (SCAN) at Stanford University for their assistance in refining the study protocol and in recruiting participants.

---

**Fig. 9.**
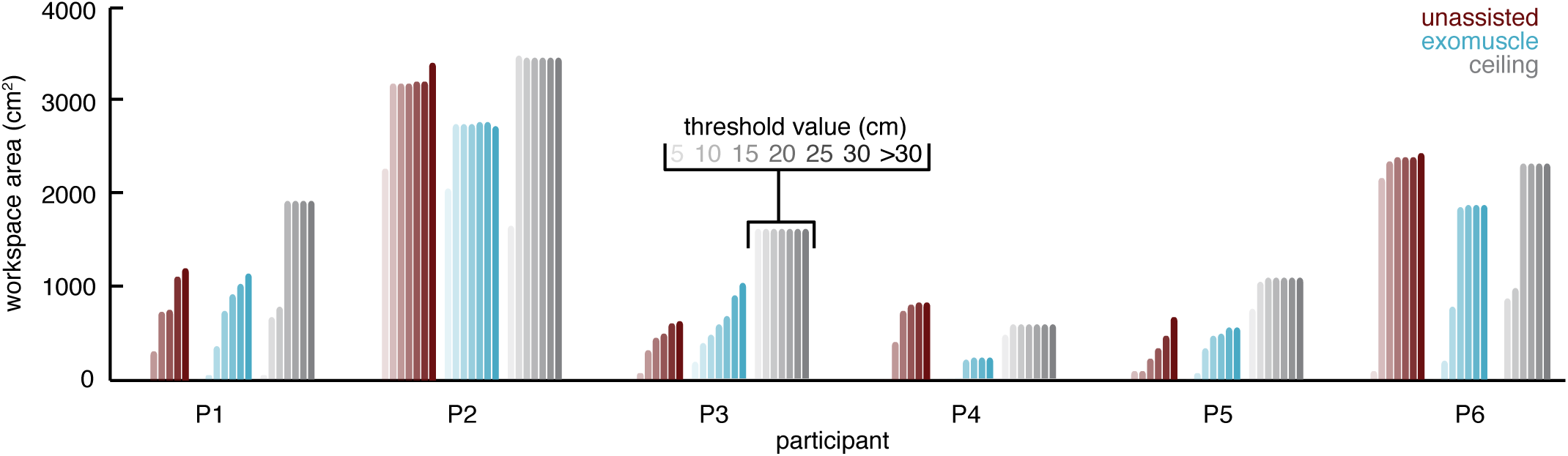
Sensitivity of workspace area measurement to the threshold value. We computed the reachable workspace area using data in which the hand and elbow were within a thresholded height of the shoulder of 20 cm. Here we compute the reachable workspace area for different threshold values ranging from 5 cm to considering all points (>30 cm) for each participant in each support condition.

